# Transcriptomics sheds light on N_2_-fixation strategies employed by a thermophilic member of the *Methanococcales*

**DOI:** 10.1101/2022.08.26.505421

**Authors:** Nevena Maslać, Chandni Sidhu, Hanno Teeling, Tristan Wagner

## Abstract

Some marine thermophilic methanogens are able to perform energy-consuming nitrogen fixation despite deriving only little energy from hydrogenotrophic methanogenesis. We studied this process in *Methanothermococcus thermolithotrophicus* DSM 2095, a methanogenic archaeon of the order *Methanococcales*, that contributes to the nitrogen pool in some marine environments. We successfully grew this archaeon under diazotrophic conditions in both batch and fermenter cultures, reaching the highest cell density reported so far. Diazotrophic growth depended strictly on molybdenum and, in contrast to other diazotrophs, was not inhibited by tungstate or vanadate. This suggests an elaborate control of metal uptake and a specific metal recognition system for the insertion into the nitrogenase cofactor. Differential transcriptomics of *M. thermolithotrophicus* grown under diazotrophic conditions with ammonium-fed cultures as controls revealed upregulation of the nitrogenase machinery including chaperones, regulators, and molybdate-importers, as well as simultaneous upregulation of an ammonium-transporter and a putative pathway for nitrate/nitrite utilization. The organism thus employs multiple synergistic strategies for uptake of nitrogen nutrients during the early exponential growth phase without altering transcription levels for genes involved in methanogenesis. As a counterpart, genes coding for transcription and translation processes were downregulated, highlighting the maintenance of an intricate metabolic balance to deal with energy constraints and nutrient limitations imposed by diazotrophy. This switch in the metabolic balance included unexpected processes, such as upregulation of the CRISPR-Cas system, probably caused by drastic changes in transcription levels of putative mobile and virus-like elements.

**Importance:** The thermophilic anaerobic archaeon, *M. thermolithotrophicus*, is a particularly suitable model organism to study the coupling of methanogenesis to diazotrophy. Likewise, its capability to simultaneously reduce N_2_/CO_2_ into NH_3_/CH_4_ with H_2_ makes it a viable target for biofuel production. We optimized *M. thermolithotrophicus* cultivation, resulting in considerably higher cell yields and enabling the successful establishment of N_2_-fixing bioreactors. Improved understanding of the N_2_-fixation process would provide novel insights into metabolic adaptations that allow this energy-limited extremophile to thrive under diazotrophy, for instance by investigating its physiology and uncharacterized nitrogenase. We demonstrate that diazotrophic growth of *M. thermolithotrophicus* is exclusively dependent on molybdenum, and complementary transcriptomics corroborated the expression of the molybdenum nitrogenase system. Further analyses of differentially expressed genes during diazotrophy across three cultivation time points revealed insights into the response to nitrogen limitation and the coordination of core metabolic processes.

## Introduction

Methanogenic archaea generate about one gigaton of methane per year (1). This amounts to about half of the greenhouse gas methane in our atmosphere (2, 3). Methanogenesis is a strictly anaerobic process occurring in habitats in which electron acceptors other than CO_2_ are depleted (4). It is accepted that under natural conditions methanogenesis provides an extremely low energy yield, causing methanogens to thrive close to the thermodynamic limits of life (1, 5, 6) Therefore, it came as a surprise when in 1984 two studies proved that *Methanothermococcus thermolithotrophicus* (7) and *Methanosarcina barkeri* (8), can perform nitrogen fixation, a very energy-demanding metabolic process that requires the hydrolysis of at least 16 ATP per molecule of fixed N_2_ (9).

The *nifH* gene is widely used as a marker to identify nitrogen-fixing organisms. Multiple environmental studies detected *nifH* genes and transcripts from methanogens in diverse anoxic habitats, such as deep seawater and hydrothermal vent fluids (10), oligotrophic open seas (11), deep-sea methane seep sediments (12), and N_2_-limited soils of salt marshes (13). Collectively these findings show that methanogens are indeed actively fixing N_2_ in nature and thereby contribute considerably to global nitrogen cycling. It also implies that methanogens and anaerobic methanotrophs (ANMEs) (14) can overcome the largest activation barrier in biology of +251 kJ.mol^-1^ (15) to break the N_2_ triple bond.

The reduction of N_2_ to NH_3_ is catalyzed by the nitrogenase enzyme complex. The overall organization of this complex is highly conserved among *Bacteria* and *Archaea*. It is composed of a dinitrogenase reductase (iron protein, NifH) and dinitrogenase (iron-molybdenum protein, NifDK) containing one [MoFe7S9C-(R)-homocitrate] iron-molybdenum cofactor (FeMo-co) and a [8Fe-7S] P-cluster (15). The nitrogenase-encoding *nif* genes and a plethora of accessory proteins with roles in regulation, biosynthesis and maturation of metal cofactors, can be clustered within one or in several operons or regulons depending on the organism (15–18). In addition to Nif, representing the most widespread and well-studied system, at least two alternative nitrogenases are known: vanadium nitrogenase (VFe protein: Vnf) and iron-only nitrogenase (FeFe protein: Anf) (15). Both are considered to be evolutionary related to the Nif system (19).

Phylogenetic analyses suggest that the ancestral nitrogenase originated in anaerobic methanogenic archaea, from where it was subsequently transferred into the bacterial domain, most probably initially to the *Firmicutes* (15, 19–22). Despite the shared origin and a minimal conserved set of genes required for functionality, archaeal nitrogenases are very distinct from their bacterial homologs. This is reflected in the genetic organization of nitrogenase operon(s), regulation of their expression by the repressor NrpR and activator NrpA, and the unique mode of post-translational activity regulation through direct protein-protein interactions (23–25). These differences have been extensively studied in the mesophilic methanogen *Methanococcus maripaludis* (16, 17, 26–30) and different *Methanosarcina* strains (31–36) using well-established genetic systems for both species. In this context it is noteworthy that *M. maripaludis* features a single operon consisting exclusively of *nif* genes (16, 28) (Fig. S1), whereas some *Methanosarcinales* harbor additional *vnf* and *anf* operons coding for V- and Fe-only nitrogenases, respectively. However, these operons all share the core organization of the *nif* operon in which *nifH*, *nifD* and *nifK* genes code for the two structural subunits of the nitrogenase system, while *nifE* and *nifN* play a putative role in the synthesis of FeMo-co. Genes *nifI_1_* and *nifI_2_* encode P_II_-family regulators that are conserved in archaea, while *nifX* has no homology with the homonymous bacterial gene (Fig. S1). NifX, absent in *Methanosarcinales*, has an unknown function. It is not required for nitrogen fixation in *M. maripaludis*, since *nifX* in-frame deletion mutants are still capable of diazotrophic growth (17).

Only a few studies exist on the diazotrophic physiology of (hyper)-thermophilic *Methanococcales*. The N_2_-fixation capabilities of these hydrogenotrophic methanogens are particularly interesting for several reasons: (*i*) their nitrogenases form a separate evolutionary branch (19) (Fig. S2, Fig. S3), (*ii*) they can perform N_2_-fixation under very high H_2_-partial pressure usually inhibiting nitrogenases (7, 8, 37), and (*iii*) they can fix nitrogen at the highest temperature ever described so far (i.e. up to 92 °C) (37). Together this suggests a unique adaptation that allows N_2_-fixation to operate at both high temperatures (7) and extreme energy limitations.

We investigated the integration of N_2_-fixation with other metabolic processes in thermophilic *M. thermolithotrophicus* by using a combined approach of physiological tests and differential transcriptomics. Our results provide new insights into the adaptive strategies of *M. thermolithotrophicus* to energy and nutrient limitation stress inflicted by Mo-dependent diazotrophy.

## Results

### Nitrogen acquisition by *M. thermolithotrophicus*

We selected *M. thermolithotrophicus* DSM 2095 due to its remarkable chemolithoautotrophic capabilities and fast growth at 65 °C in mineral medium (38, 39). Series of incubations with different NH_4_Cl concentrations, ranging from 0.1 to 16.8 mM, showed that *M. thermolithotrophicus* required a minimum of 10 mM NH_4_Cl for best growth (Fig. 1A) in our optimized medium (see Materials and Methods for composition) (39, 40). Higher NH_4_Cl concentrations of 25-200 mM NH_4_Cl did not result in higher cell yields, and a 500 mM excess of NH_4_Cl led to a decrease (Fig. S4). Continuous monitoring of ammonia consumption during growth on 10 mM NH_4_Cl over a span of 26 hours confirmed depletion proportional to the observed increase in biomass (Fig. 1B). The culture reached stationary phase after 18 h during which most of the ammonia was consumed beyond the reliable detection limit.

**FIG. 1.**
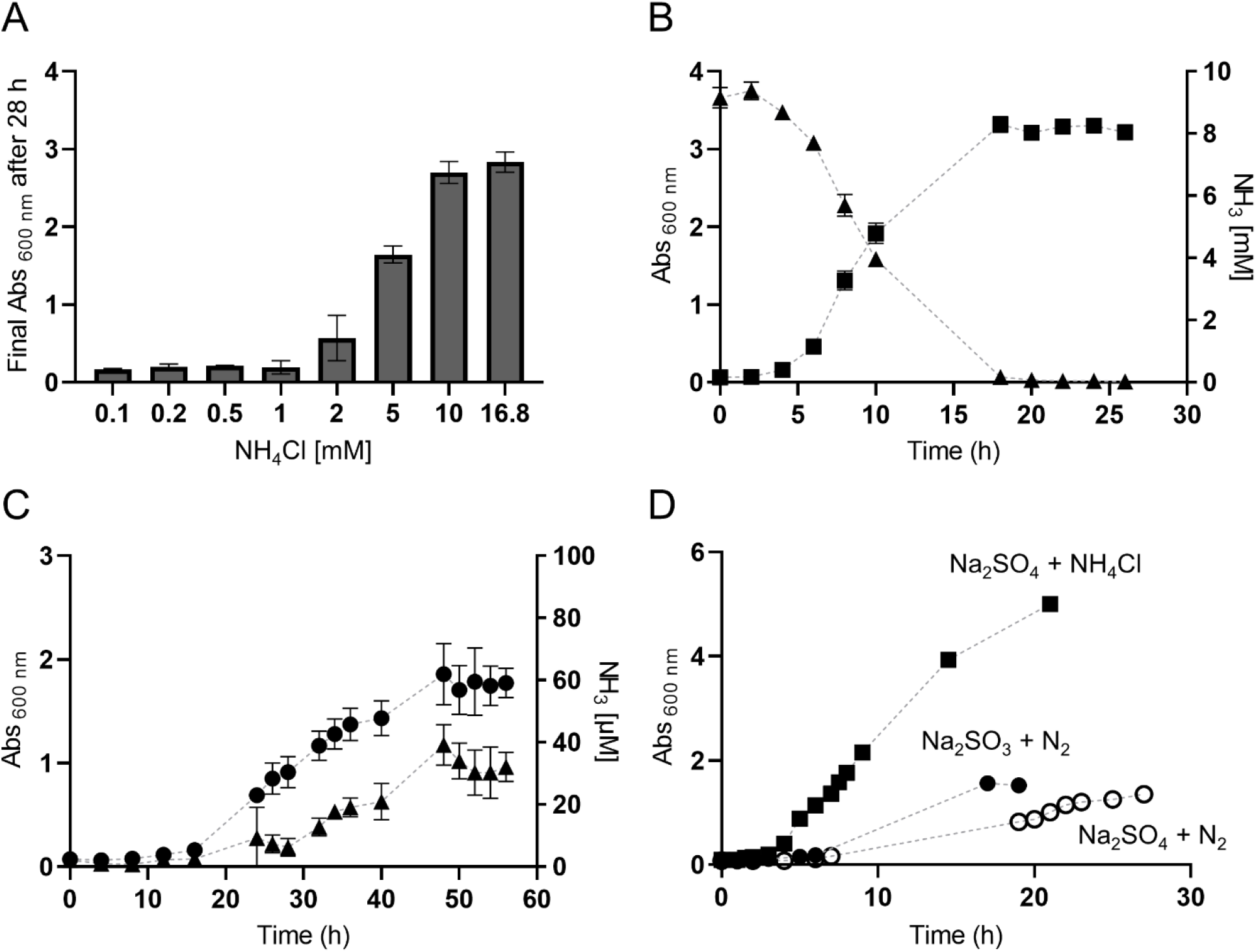
Growth of *Methanothermococcus thermolithotrophicus* on different nitrogen sources. (A) Final Abs _600 nm_ of *M. thermolithotrophicus* cultures after 28 h of incubation with different NH_4_Cl concentrations. (B) Growth curve of *M. thermolithotrophicus* cultures grown on 10 mM NH_4_Cl (squares) and NH_4_Cl consumption during growth (triangles). (C) Growth curve of diazotrophic *M. thermolithotrophicus* cultures grown on N_2_ as the sole nitrogen source (circles) and NH_3_ release during the growth (triangles). (D) Growth curves of non-diazotrophic *M. thermolithotrophicus* on 16.8 mM NH_4_Cl (squares), a diazotrophic culture grown on Na_2_SO_3_ (full circles) and a diazotrophic culture grown on Na_2_SO_4_ (empty circles) in a fermenter. Measurements for A have been performed in duplicates. Measurements for B and C have been performed in triplicates.

*M. thermolithotrophicus* was adapted to diazotrophic conditions after three successive transfers to NH_4_Cl-free medium. Under this condition, dissolved N_2_ was the sole available nitrogen source. The cell density of the diazotrophic culture was six times higher than reported by Belay and colleagues in their original study on the discovery of diazotrophy in methanogens in 1984 (e.g., 1.85 achieved in 48 h as compared to 0.4 achieved in ~20 h (7)). Ammonia possibly released in the medium was measured during diazotrophic growth (Fig. 1C), and traces were detected during the mid-exponential phase reaching a maximum of 40 μM ammonia in the stationary phase (Fig. 1C). However, we cannot exclude that detected ammonia also originated from cell lysis rather than from active excretion.

In addition to batch diazotrophic cultures, *M. thermolithotrophicus* was successfully grown in a 10 L fermenter continuously supplied with H_2_/CO_2_ and N_2_. To maintain the sulfur source in the medium, we replaced Na_2_S used in batch cultures (which would be flushed out as H_2_S) with Na_2_SO_3_ or Na_2_SO_4_ (see Materials and Methods)(39). Diazotrophic growth was not affected by HSO_3_^-^, a known inhibitor of methanogenesis (Fig. 1D) (41). While the final cell yield was similar to the one observed in batch cultures, division times were shorter (Fig. 1D). Fermenter-grown cultures supplemented with NH_4_Cl had higher final yields than diazotrophic fermenter-grown cultures. However, in comparison to NH_4_^+^-grown batch cultures, the difference was not very pronounced, as final yields were only slightly higher and the division times were similar.

### N_2_-fixation is molybdenum-dependent

We then investigated the nitrogenase type used for diazotrophy, taking into account that the *M. thermolithotrophicus* genome features only one *nifDK*. It must be noted that *M. thermolithotrophicus* resembles *M. maripaludis* in terms of physiology, and the latter was shown to require molybdenum (Mo) for its diazotrophic growth (28). Therefore, we first tested for molybdenum (Mo) dependency. Growth of cultures lacking either Mo, vanadium (V) or both metals in the medium were monitored simultaneously. Depletion of Mo and V from the media was achieved by three successive culture transfers to the same media without Mo and V. The highest optical density (OD_600 nm_) was reached when *M. thermolithotrophicus* was grown in the control medium containing both Mo and V. Growth was similar in the diazotrophic culture incubated without V, but with a lower final OD_600 nm_ (Fig. 2A). This trend was reproducible and suggests that V acts as a potential growth stimulator under such conditions. No growth was observed in the absence of Mo, with or without V (Fig. 2A). The minimal required Mo concentration of 0.1 μM was determined in a separate series of incubations with Mo concentrations ranging from 0.01 to 100 μM (Fig. 2B). To check, if Mo is essential for methanogenesis (42) or metabolic processes other than nitrogen fixation, we supplemented the culture grown in absence of Mo with NH_4_Cl, which restored growth after an overnight incubation (Fig. 2A).

**FIG. 2.**
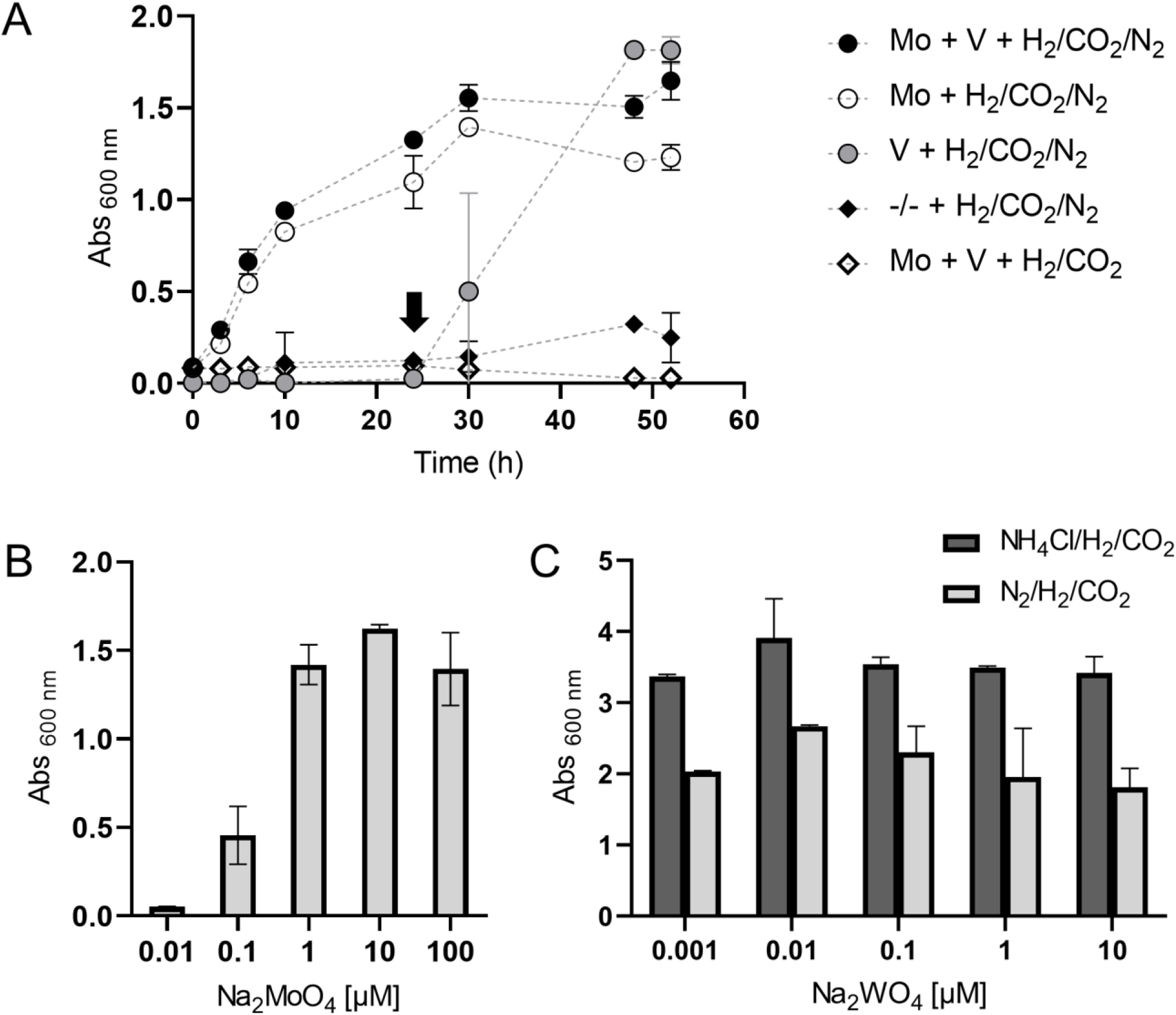
Influence of trace metal availability on diazotrophic growth of *M. thermolithotrophicus*. (A) Growth curves of diazotrophic *M. thermolithotrophicus* grown in the medium with both Mo and V (full circles), without V (empty circles), without Mo (gray circles), without both Mo and V (full rhomboids) and negative control (empty rhomboids). The arrow indicates the supplementation of the culture without Mo with 16.8 mM NH_4_Cl. (B) Final Abs _600 nm_ of *M. thermolithotrophicus* as a function of [MoO_4_^2+^] concentration. (C) Final Abs _600 nm_ of non-diazotrophic (dark gray bars) and diazotrophic (light gray bars) *M. thermolithotrophicus* as a function of [WO_4_^2+^] concentration. 10 μM of Na_2_MoO_4_ was used for this experiment. Measurements for A have been performed in triplicates. Measurements for B and C have been performed in duplicates.

An inhibitory effect of tungstate (WO_4_^2-^) on diazotrophic growth has already been observed in *A. vinelandii* OP (43), *Methanosarcina barkeri* 227 (32) and *M. maripaludis* strain S2 (28). In this case, WO_4_^2-^ can inhibit MoO_4_^2-^ uptake (44) and it can be incorporated into the nitrogenase, thereby rendering the protein inactive (43, 45, 46). These inhibition mechanisms are competitive and thus dependent on the MoO_4_^2-^/WO_4_^2-^ ratio. Surprisingly, there was no observable inhibitory effect in *M. thermolithotrophicus* (Fig. 2C), even at a 1:1 MoO_4_^2-^/WO_4_^2-^ ratio. In contrast, diazotrophic growth of *M. maripaludis* was inhibited at a 10:1 MoO_4_^2-^/WO_4_^2-^ ratio (28). This observation might be explained by different affinities of the WO_4_^2-^ and MoO_4_^2-^ transport systems in both species. The ModABC transporter (47, 48) is present in both organisms and should transport MoO_4_^2-^ and WO_4_^2-^ (49). In addition, *M. thermolithotrophicus* has a highly specific WO_4_^2-^ transporter TupABC (50) that is lacking in *M. maripaludis*. Instead, *M. maripaludis* has a third type of tungstate transporter WtpABC (49, 51), which can transport WO_4_^2-^ and Mo_4_^2-^ but has a higher affinity for WO_4_^2-^ than has been shown for Mod and Tup in *Pyrococcus furiosus* (51). Furthermore, it is also known that Mod transporters can have different affinities for both oxyanions in different organisms (49, 52, 53), suggesting that both transporter specificities and the specificity of the FeMo-co insertion machinery contribute to the W tolerance during diazotrophy.

### Diazotrophy shifts expression of a large number of genes

Transcriptomics and DNA-microarrays have been used to investigate the complex metabolic and regulatory networks that control N_2_-fixation in two model organisms*: Azotobacter vinelandii* (54) and *Methanosarcina mazei* Gö1 (36, 55, 56). These studies led to the discovery of three gene clusters (*rnf1, rnf2* and *fix*) coding for electron transfer systems that provide reducing equivalents to the nitrogenase in *A. vinelandii*, as well as multiple potential transcriptional and sRNA regulators in *Methanosarcina mazei* Gö1. Here, we used comparative transcriptome profiling of N_2_-fixing *M. thermolithotrophicus* cultures versus NH_4_^+^-grown cultures to investigate the metabolic adaptations induced by nitrogen fixation. The experiment was conducted in biological triplicates at three different time points: 3 h (early exponential phase), 21 h (stationary phase-starvation due to the exhaustion of the gas phase), and 25 h post-inoculation (see Materials and Methods, Fig. 3A, and Supplementary Table 1). The gas phase was exchanged after sampling at the 21 h time point, since the cultures consumed H_2_/CO_2_. The robustness of our sample data was reflected in a corresponding principal component analysis (PCA), in which triplicates from different conditions and time points are clustered (Fig. 3B). More importantly, it revealed that 92% of the variation could be explained by the different sample treatments.

**FIG. 3.**
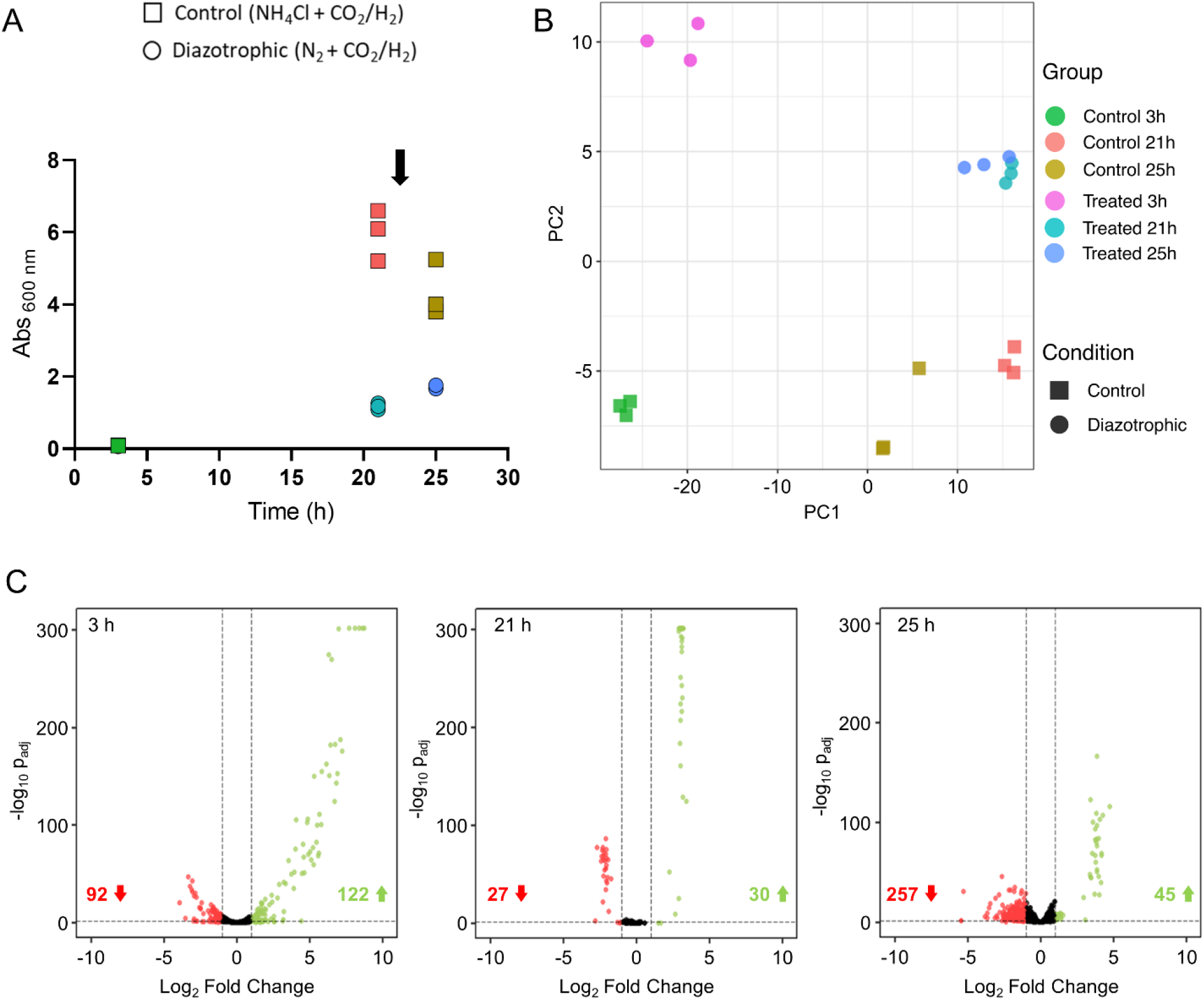
Differential gene expression between NH_4_Cl-grown and diazotrophic cultures. (A) Points during growth of the control (squares) and diazotrophic (circles) cultures at which samples for transcriptomic profiling were taken (each of the triplicates is shown separately). The arrow indicates the complete gas phase exchange of the cultures after the sample at 21 h was taken. (B) PCA plot showing the first two principal components that explain the variability in the data using the regularized log count data. Control corresponds to the NH_4_Cl grown cultures. (C) Volcano plots of differentially expressed genes in the diazotrophic culture versus control at different time points. Negative and positive log_2_ fold changes >1 with adjusted p < 0.05 are shown in red and green, respectively. Values in red and green indicate the number of down- and up-regulated genes, respectively.

Transcriptome profiling across the three time-points revealed prominent changes (Fig. 3C). Out of the total 1,751 predicted genes (open reading frames, ORFs), 1,737 genes were found to have non-zero read counts. At 3 h, 12.3% (214) were differentially expressed with most (122/214) being upregulated under diazotrophic conditions. More than half of these genes (70/122) were of an unknown function without any homologs in the databases (their probable role is discussed further below). The difference was smaller at 21 h, with only 3.3% (57) of the total expressed genes having differential expressions, however at 25 h, 17.2% (302) were differentially expressed, with most of them downregulated (257/302) under diazotrophic conditions (Supplementary Table 2).

To check the overall expression of genes irrespective of differential expression, we mapped reads to predicted genes in terms of transcripts per million (TPM). Details of this analysis are summarized in the sections below.

### Nitrogen- and molybdate-acquisition genes are mostly affected during the early exponential phase

As anticipated, upregulated genes at 3 h included the *nif* operon (*nifHI1I2DKXEN*) and molybdate-acquisition genes (*modABC*). This illustrates a swift response of *M. thermolithotrophicus* to diazotrophic conditions. The highest log_2_-fold change (FC) was observed for *nifI_1_I_2_HDKX* genes with values ranging from 8.8 to 7.0 (Supplementary Table 2). The proteins of the FeMo-co biosynthetic machinery, *nifE and nifN*, were also among the upregulated genes with log_2_FC of 5.3 and 4.6 respectively (Fig. 4A-B). An essential gene for FeMo-co biosynthesis, *nifB*, (57, 58), was transcribed at a lower level with a log_2_FC of 1.8.

**FIG. 4.**
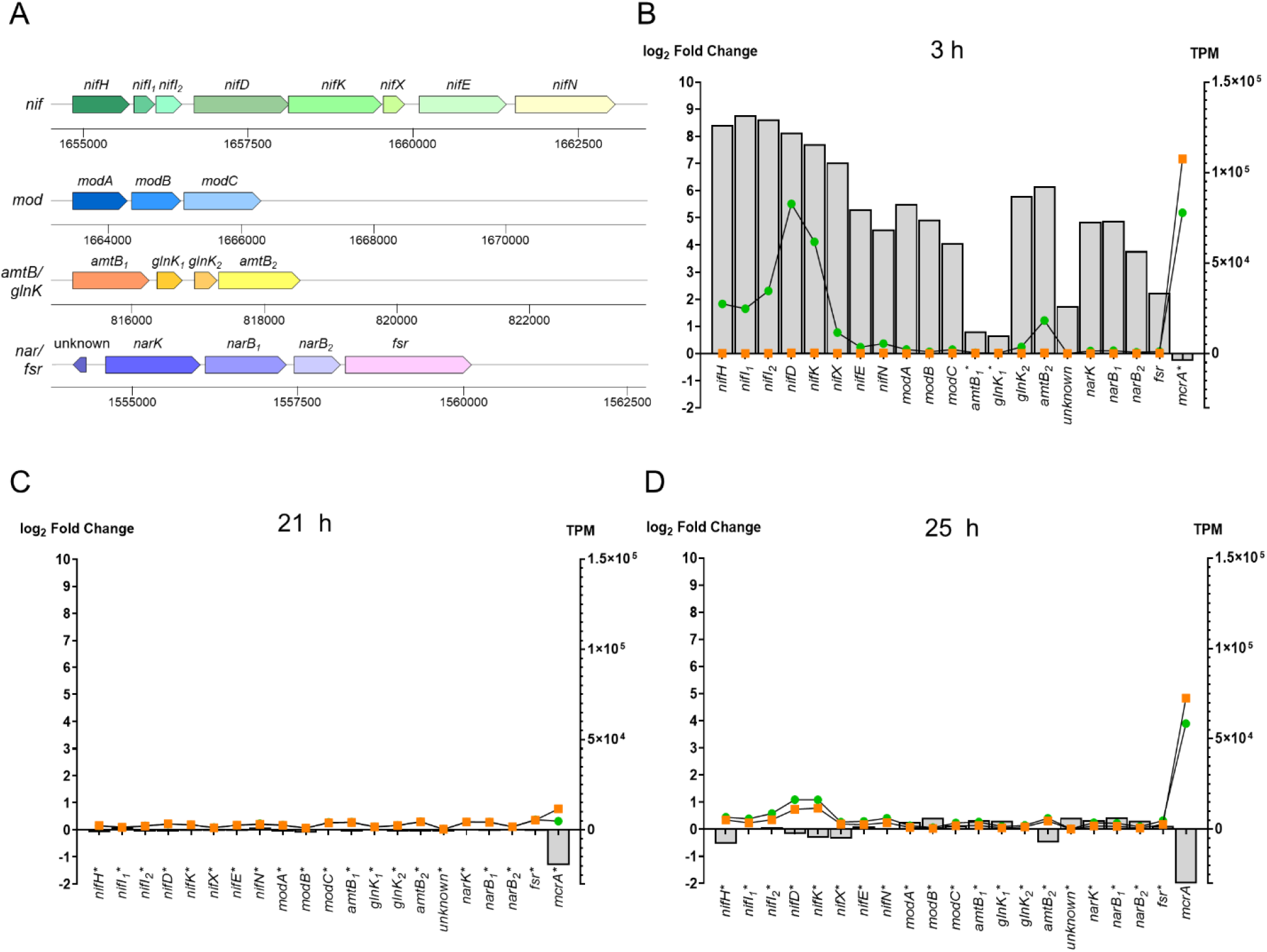
Changes in the expression over time of selected genes involved in nitrogen acquisition. (A) Arrangement of the *nif, mod, amtB/glnK* and *nar/fsr* operons in *M. thermolithotrophicus*. (B-D) Log_2_-fold changes (light gray bars) and transcript numbers per million (TPM) of the controls (orange squares) and diazotrophic cultures (green circles) of nitrogen acquisition genes compared tothe *mcrA* gene as reference after 3 h (B), 21 h (C) and 25 h (D). Genes that were not differentially expressed at a given time point are marked by an asterisk.

The *mod* operon, coding for the three subunits of a molybdate ABC transporter (*modABC*), was also upregulated (Fig. 4A), with *modA* being strongly expressed with a log_2_FC of 5.5 (Supplementary Table 2). This agrees with our physiological data that show a clear Modependency under diazotrophic conditions. Although three *modA* instances are present in the genome, only the *modA* gene in vicinity of *nif* operon that is part of the complete *mod* operon was highly expressed. The *tupA* gene coding for the tungstate transporter was also upregulated (log_2_FC: 1.7), strengthening the hypothesis that both transporter specificities and adaptation of the cofactor biosynthetic machinery contribute to the W tolerance during diazotrophy.

The other genes upregulated at 3 h included the ammonium transporter *amtB_2_* and its P_II_-family regulatory protein *glnK_2_* with the log_2_FC of 6.2 and 5.8, respectively (Supplementary Table 2). The genome of *M. thermolithotrophicus* harbors two different *amtB* genes with their associated *glnK* regulators (40). Interestingly, *amtB_1_* and *glnK_1_* were not differentially expressed (Supplementary Table 2, Figure 4A-B), which might point to the existence of an internal regulator or an additional promotor. In addition to the *nif* and *amtB/glnK* operons, which are known to be upregulated during diazotrophy from previous studies (54, 55, 59), the ammonium assimilating glutamine synthetase (*glnA* gene was also upregulated (Supplementary Table 2). However, the transcript level of the second enzyme involved in the N-assimilation, the glutamate synthase, remained unchanged, suggesting that GlnA is the rate limiting step in ammonium assimilation.

Unexpectedly, *narK* coding for a putative nitrate transporter and *narB* coding for a molybdopterin-dependent nitrate reductase (60), were also highly expressed during diazotrophy in the early exponential stage (Fig. 4A-B, Supplementary Table 2). If nitrate would be imported and reduced in the cell, the oxidant nitrite would be generated and could damage the highly oxidation-sensitive methanogenic machinery (61). A gene coding for a F420-sulfite reductase isoform (also belonging to Fsr group 1 but different to the one naturally expressed under sulfite condition described in (39)), which co-occurs with *narK* and *narB*, would be a plausible candidate for nitrite detoxification, since Fsr has been recognized to catalyze nitrite reduction (39). *M. thermolithotrophicus* has been reported to be able to grow on nitrate as the only source of nitrogen (62), which might involve these genes.

In addition, three genes coding for the enzymes of the molybdopterin biosynthetic pathway (*moaE, mobB* and *moeB*) were upregulated. They might be involved in supplying the putative nitrate reductase with molybdopterin, while the formylmethanofuran dehydrogenase enzyme used for methanogenesis harbors a tungstopterin (42). The pathway of molybdopterin biosynthesis has been extensively studied and it is highly conserved (63). The pathway of tungstopterin biosynthesis is thought to be homologous up to the step of metal insertion (64). It is proposed that the biosynthetic machinery is able to distinguish between the two metals and insert the correct metal into the respective enzymes by employing MoeA isoenzymes selective for either molybdate or tungstate (64, 65). All archaeal genomes sequenced so far encode two *moeA* isoforms sharing around 40% identity (64). This might explain how organisms are able to correctly express different proteins with molybdopterin and tungstopterin simultaneously (64). However, this hypothesis still lacks experimental validation.

As described in Fig. 4C-D, the expression of genes partaking in N_2_-fixation and molybdenum acquisition remained unchanged at 21 h and 25 h and therefore, no differential expression was observed. The overall expression of genes in terms of TPM followed a similar pattern. An established marker for metabolic activity of methanogens, the transcription level of *mcrA* (66), was used as a reference point for comparison (Fig. 4B-D).

None of the methanogenesis pathway genes showed any difference in transcription levels at the early exponential phase. However, after 25 h nearly all genes involved in methanogenesis were downregulated. The expression of F420-reducing hydrogenase was maintained at the same level at all time-points to supply reduced F420 from H_2_-oxidation. Hydrogenotrophic *Methanococcales* can alternatively use formate as electron source (67, 68), and in this case a putative formate transporter (*fdhC*) and formate dehydrogenase (*fdhF*) were found to be downregulated at 3 h. In addition to methanogenesis, numerous anabolic processes were shut down at 25 h, such as carbon-assimilation (e.g. pyruvate:ferredoxin oxidoreductase), amino-acid metabolism (e.g. ketol-acid reductoisomerase), lipid biosynthesis (e.g. hydroxymethylglutaryl-CoA synthase), ATP synthesis (i.e. ATP-synthase) and vitamins and coenzymes biosynthesis (e.g. *hemE*). Such a decrease in catabolic and anabolic processes combined with the downregulation of genes involved in S-layer formation and cellular division (e.g. *ftsZ*) suggests a fine-tuned metabolic mode of energy saving to prioritize nitrogen fixation.

### Transcriptional and translational machineries are considerably downregulated

Both the transcriptional and translational machineries responded negatively to diazotrophic conditions after 3 h. This included downregulation of RNA polymerase subunits (*rpoA2H*), the sigma factor 70 (*rpoD*, controlling the transcription of housekeeping genes), and 35 ribosomal proteins (Supplementary Table 2). Our ribosomal RNA expression data has the limitation that rRNA removal treatment was performed before sequencing. Still, we could detect downregulation of rRNA expression, which was corroborated by downregulation of ribosomal proteins. Some genes with a putative function in tRNA/ribosome biogenesis and biosynthesis of nucleotide and amino acid precursors were also negatively affected, corroborating a deep impact on the overall translation process under N_2_-fixing conditions. For example, two key enzymes (transketolase and transaldolase) of the non-oxidative branch of the pentose phosphate pathway (69) for synthesis of nucleotides and histidine (from ribose 5-phosphate), as well as aromatic amino acids (from erythrose 4-phosphate) precursors were downregulated (Supplementary Table 2). This is another notable difference from *Methanosarcina*, in which the upregulation of genes involved in the synthesis of aromatic amino acids was observed (55). Notably, tRNA^Thr^ and tRNA^Ser^ were upregulated while tRNA^Ala^ and tRNA^Pro^ were downregulated at 3 h. Taken together, changes in the expression of all the mentioned genes contribute to the restriction of the entire translation process. The DNA replication system seems not to be impacted, since we could not observe any changes in the transcript levels of DNA polymerase encoding genes. This is in line with the results of Belay and coworkers, who showed that during diazotrophic growth the cellular protein content of *M. thermolithotrophicus* was significantly reduced (7).

### High expression of CRISPR-Cas genes and putative viral genes

Components of the CRISPR-Cas virus defense system were also upregulated under diazotrophy at 3 h, including Csm1-5 of the Csm effector complex and CRISPR-associated proteins (Cas proteins) Cas4-6 (Supplementary Table 2). This might be explained by presence of a prophage that is expressed when the cells are energy depleted or otherwise stressed. The Phaster server, a tool for finding prophages in bacterial genomes (70), was unsuccessful in detecting any complete prophages in the *M. thermolithotrophicus* genome. However, we identified a locus of 28 co-occurring ORFs that were highly expressed at all-time points (Fig. 5). Twenty out of these genes had no characterized homologs, while eight encoded putative virus-like, replication, and mobile genetic elements. We used models generated by Alphafold2 (71) and a membrane prediction tool to gain further information on these sequences and examined 18 confident models (Fig. 5). Five of these proteins were predicted to be secreted or embedded in the membrane, but structural homologs were scarce based on the predicted models (Supplementary Table 3). The protein encoded by the third gene of the locus (Fig. 5) has already been structurally characterized in *Thermococcales* (72) and is believed to be a virus-like element. While *Methanococcales* share some of these genes, *Methanosarcina mazei*, a model organism for the *Methanosarcinales*, contains only two of them in its genome (Supplementary Table 3). These 28 co-occurring genes might be derived from a small plasmid transferred by conjugation, or represent a yet unknown prophage, although we could not identify a protein that could be responsible for independent insertion or replication of this element. This region could therefore be considered part of the dark matter in archaeal genomes (73). However, active transcription of all the genes of this locus might be a plausible explanation for the observed upregulation of the CRISPR-Cas system.

**FIG. 5.**
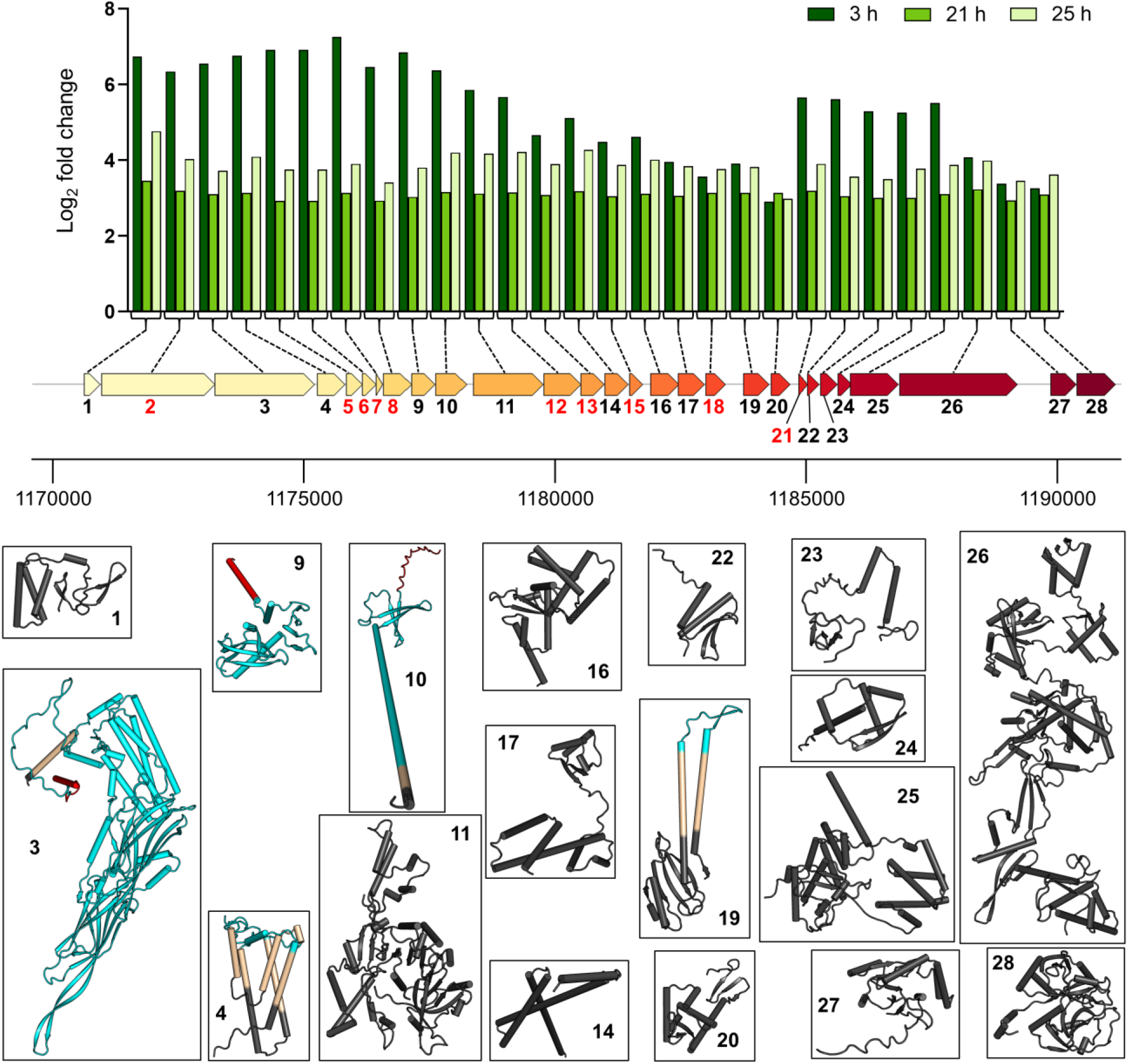
Log_2_ fold changes over time for 28 co-occurring genes of unknown function that are of putative viral origin. AlphaFold 2 (72) models are represented by cartoons and color-coded as follows: black for intracellular segments, cyan for predicted extracellular segments, ocher for transmembrane segments and red for predicted signal sequences. The topology prediction was made via the DeepTMHMM server (106). Genes with numbered red labels highlight models with an overall per-residue confidence score (pLDDT) below 75. These models are not presented due to their low confidence scores.

Another locus of 26 co-occurring ORFs was downregulated under diazotrophic conditions at all-time points (Fig. S5). Again, most of these genes had unknown functions, with the exception of a putative transcriptional regulator, a mini chromosome maintenance protein, and a recombinase (Supplementary Table 4). Four of these genes were predicted to encode secreted proteins, and eight contained transmembrane segments. Based on these observations, we assume that these 26 co-occurring open reading frames are also derived from mobile elements or prophage-associated genes.

## Discussion

Diazotrophy allows microbes to survive when nitrogen becomes limited in their natural habitats. This is also the case for (hyper)thermophilic methanogens, such as *M. thermolithotrophicus*, which has been shown to rely on diazotrophy in different environments (10–11, 13). A notable exception among *Methanococcales* is *Methanocaldococcus jannaschii*, a non-diazotrophic methanogen isolated from a white smoker on the East Pacific Rise. While ammonium concentrations at the *M. jannaschii* isolation site were not measured (74), some parts of the East Pacific Rise, such as Guaymas Basin, are known to feature notable ammonium concentrations (e.g. 15.3 mM (75)). Availability of this inorganic nitrogen source might have resulted in a complete loss of nitrogen fixing abilities in *M. jannaschii*.

Due to its active N_2_-fixation, *M. thermolithotrophicus* represents one of the contributors to the available nitrogen pool in specific environments (10–11, 13). In our laboratory cultures, *M. thermolithotrophicus* released only minute amounts of up to 40 μM ammonia to the medium, which might have just resulted from passive diffusion or cell lysis. It therefore seems that N_2_-fixation *in M. thermolithotrophicus* is precisely controlled to avoid any losses, likely by the NifI_1,2_ regulation system (Fig. 6). The excessive energy cost inflicted by nitrogenase activity was noticeable in our physiology experiments (Fig. 1), as it reduced final yields by a factor of two in both batch and fermenter grown cultures.

**FIG. 6.**
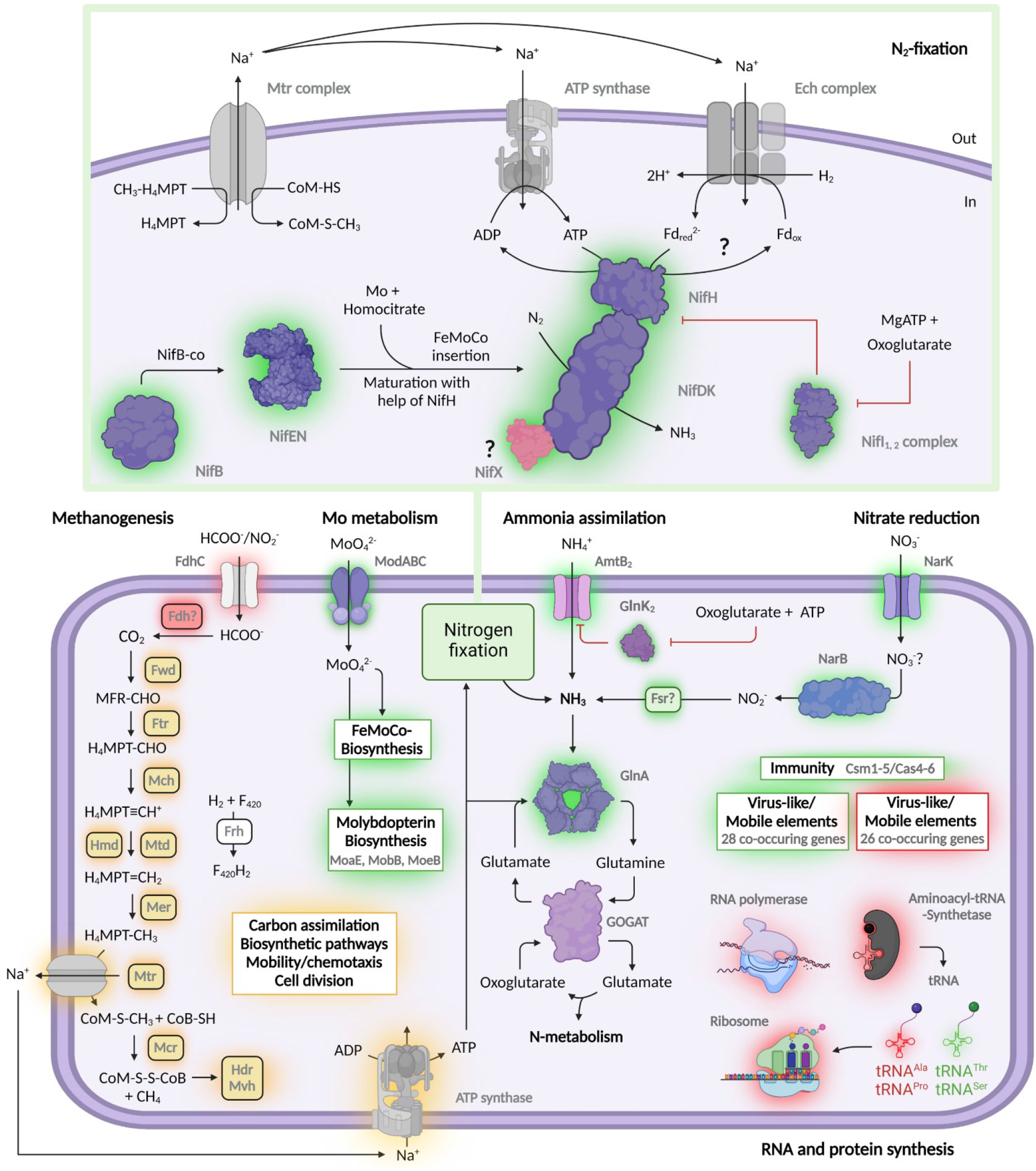
Influence of nitrogen fixation on the metabolism of *M. thermolithotrophicus*. In the scheme, differentially expressed genes involved in methanogenesis, nitrogen fixation, ammonia assimilation, nitrate reduction, molybdate import, molybdopterin biosynthesis, immunity, mobile elements, transcription and translation, as well the tRNAs are shown. Differentially upregulated and downregulated transcripts at 3 h post-inoculation are highlighted with a green and red glow, respectively. An orange glow highlights differentially downregulated gene transcripts at 25 h post-inoculation. Full names of enzymes can be found in Supplementary Table 2. This picture has been created with Biorender.

Transcriptome profiling under diazotrophic conditions revealed that *M. thermolithotrophicus* relies on multiple synergistic strategies that ensure both sufficient N_2_-fixation and energy preservation to support cellular growth (Fig. 6). In addition, *M. thermolithotrophicus* enhances nitrogen acquisition by increasing ammonia uptake via its *amtB_2_* transporter and glutamine synthetase overall activity - a strategy that has been described before based on proteomics of nitrogen-starved *M. maripaludis* cultures (59). While upregulation of the *nif* and *amtB/glnK* operons under nitrogen limitation have been previously reported in other diazotrophs (54, 55, 59), the upregulation of a putative nitrate transporter (*narK*), a nitrate reductase (*narB*), and a new isoform of F420-dependent sulfite reductase (*fsr*) reported in this study is so far unique to *M. thermolithotrophicus*.

Like *M. thermolithotrophicus*, the hyperthermophile *Methanocaldococcus infernus* has also been reported to grow on nitrate as sole nitrogen source (76). *M. infernus* was isolated from a smoking crater of the Logatchev hydrothermal vent field (76, 77) from which other nitrate-reducing organisms have been isolated as well (78). Although hydrothermal fluids have been reported to be depleted in nitrate and nitrite (79), bottom seawater can contain nitrate for use as electron acceptor and nitrogen source. The enzyme which can possibly be used for intracellular nitrate reduction by these methanogens is the nitrate reductase NarB. NarB is expected to harbor a tungstopterin or molybdopterin cofactor, which might explain upregulation of molybdopterin biosynthesis genes (*moaE*, *mobB*, *moeB*) under diazotrophic conditions. In this context, the molybdate transporter Mod would play a dual role, supplying Mo for both the nitrogenase and NarB metallo-cofactor.

Strict dependence on Mo for diazotrophy has also been described for *Methanococcus maripaludis* (28). Likewise, our results support that the single nitrogenase operon of *M. thermolithotrophicus* encodes a molybdenum nitrogenase. This is corroborated by the *M. thermolithotrophicus* phylogenetic position (19), as well as high transcription levels of the *mod* operon (encoding a molybdate ABC transporter) adjacent to the nitrogenase operon under diazotrophy. In addition to *modA* within the *mod* operon, *M. thermolithotrophicus* features two additional *modA* instances. These are not co-localized with *modBC* genes and were not differentially expressed in our experiments. Although multiple *modA* genes are present in different diazotrophic organisms, including *A. vinelandii* and the methanogens *M. maripaludis* and *M. mazei*, the benefit of multiple *modA* genes is not understood yet. Interestingly, in *M. mazei* (55), no upregulation of Mod transporters was observed. In *Methanosarcinales* expression of *modABC* is under the control of the transcriptional regulator ModE. This regulator also affects transcriptional regulation of different nitrogenases types, if these are present (80, 81). *Methanococcales* are devoid of *modE* homologs, which might explain the observed upregulation of the *mod* operon under diazotrophic growth in *M. thermolithotrophicus*.

The *nif* operon exhibited the highest log_2_FC values between both conditions. However, changes in transcript levels of the *nifE* and *nifN* FeMo-co biosynthesis genes were less prominent than those of the structural *nif* genes. The former are possibly required in lower amounts, as was the case for the FeMo-co biosynthesis gene *nifB*. Levels of *nifX* transcription were similar to those of the *nifDK* nitrogenase genes, suggesting potential association with the nitrogenase complex. The *nifX* gene is not essential for nitrogen fixation in *M. maripaludis*, and has not been detected in the nitrogenase complex of this archaeon. Instead, it has been proposed to take on the role of the FeMo-co biosynthesis genes (17). However, since *M. maripaludis nifX* is not homologous to *M. thermolithotrophicus nifX*, future experimental evidence is required to clarify its putative function and association to NifDK in (hyper)-thermophilic methanogens.

The enormous ATP investment required for diazotrophy did not affect the transcription of methanogenesis genes at the onset of diazotrophic growth. This observation corroborates previous studies (55, 59) stating that the proportion of the overall energy required for maintenance increases when switching to diazotrophy (82). The high energy demand for nitrogen fixation is counterbalanced by drastic downregulation of the transcription and translation machineries. The question is, whether this is an *a priori* and thus targeted metabolic adaptation, or a mere general consequence of energy starvation. It has been described that transcription and translation are reduced upon nutrient limitations. After all, 40-70% of the cellular ATP pool in growing bacteria is attributed to protein synthesis (83). It is also known that bacteria can balance tRNA abundances when under stress to selectively regulate the translation of stress-induced proteins. These proteins contribute to the response and adaptation to different types of stress, including nutrient limitation (84). It has recently been shown that *M. maripaludis* adopts a resource relocation strategy upon energy depletion (85): instead of reducing ribosome numbers, the cells rather redistribute available energy and decrease catabolic and ribosomal activities; a strategy also described in bacteria (86). Therefore, our results suggest a targeted response associated with switching to a diazotrophic lifestyle. In comparison, no such changes in the transcript levels of transcriptional and translational genes were detected in either *Azotobacter vinelandii* (54) or *M. mazei* (55) when switching to diazotrophic growth, even though in the case of the former 30% of the genes is differentially expressed (as observed by differential transcriptomics), and in case of the latter 5% (as observed by microarray analysis).

Studies in *M. mazei* and *M. maripaludis* confirmed the prominent regulation of nitrogen metabolism at the transcriptional level, particularly the importance of the global nitrogen regulatory repressor NrpR of nitrogen assimilation genes (24, 25, 35, 87). NrpR represses transcription by binding to the *nif* and *glnA* promoter regions in a 2-oxoglutarate-dependent manner. NrpR from *M. thermolithotrophicus* shares 38.2% identity with the one from *M. mazei*. Interestingly, NrpR expression levels are not regulated by nitrogen supply (24, 25). Thus, although a homolog of NrpR does exist in *M. thermolithotrophicus*, its transcription level remained unchanged during diazotrophic growth. In contrast to NrpR, NrpA, the *nif* promotor-specific activator, sharing 32% identity with the one from *M. mazei*, is known to be upregulated upon nitrogen limitation (88). We did not observe this in our experiments suggesting that NrpA could be constitutively expressed in *M. thermolithotrophicus*.

One major difference between *M. mazei* and *M. thermolithotrophicus*, however is the role of regulatory sRNAs. *Methanosarcina mazei* Gö1 expresses multiple sRNA genes in response to nitrogen limitation (56), with sRNA_154_ being the best characterized (36). In *M. mazei* sRNA_154_ stabilizes the nitrogenase, *glnA_1_* and *nrpA* mRNAs (36). While sRNA_154_ is highly conserved within *Methanosarcinales*, there are no sRNA_154_ homologs in *Methanococcales* (89). Consequently we did not detect such a pattern in *M*. *thermolithotrophicus*.

Transcriptome analysis can provide valuable information on proteins involved in nitrogen-fixation. For instance, transcriptome analysis allowed the discovery that in *A. vinelandii* electron bifurcation is coupled to a flavodoxin that fuels the nitrogenase (54). However, we could not detect any putative novel candidates that could shuttle electrons to the nitrogenase, and we propose reduced ferredoxin as a candidate for electron delivery, a known electron carrier for anabolic reactions. *M. thermolithotrophicus* would use H_2_-oxidation for ferredoxin reduction, an endergonic process that requires the coupling of the influx of sodium ions by the Ech complex. The reduced ferredoxin would then drive the reduction of NifH for N_2_-fixation (Figure 6).

Here we present novel insights into the metabolic rebalancing that methanogens employ to accommodate Mo-dependent diazotrophy. Further studies at the protein level are required to decipher the mechanistic of this complex adaptation in greater detail. In particular molecular investigations of the regulatory functions of NifI_1,2_, the role of NifX, and the intrinsic properties of the thermostable NifHDK complex will unveil the secrets of the astonishing diazotrophic capabilities of *M. thermolithotrophicus*.

## Material and methods

### Growth conditions

*Methanothermococcus thermolithotrophicus* DSM 2095 (Leibniz Institute DSMZ - German Collection of Microorganisms and Cell Cultures, Braunschweig, Germany) was grown under anoxic conditions in minimal mineral media with 1 bar overpressure of either H_2_:CO_2_ (80%:20%) for non-diazotrophic or 1.2 bar overpressure H_2_:CO_2_:N_2_ (58.2%:14.5%:27.3%) for diazotrophic cultures. Cultures were grown in 250 mL serum flasks (Glasgerätebau Ochs, Bovenden, Germany) sealed with rubber stoppers and aluminium crimps in final volumes of 10 mL with a 1:10 inoculum. Serum flasks and media were made anoxic prior to inoculation by sparging with N_2_ and two final gas exchanges with H_2_:CO_2_ (80%:20%). Incubation was done at 65 °C, in the dark, without shaking.

### Medium composition

The used minimal mineral medium was prepared as described in Müller *et al*. 2021 (40), but with replacement at equal final concentration of Fe(NH_4_)_2_(SO_4_)2 x 12H_2_O by FeCl_2_ x 4H_2_O, and of Na_2_SeO_3_ x 5H_2_O by Na_2_SeO_4_. The used trace metal solution 100 fold concentrated contained 7.1 mM nitrilotriacetic acid, 0.45 mM MnCl_2_ x 2H_2_O, 0.68 mM FeCl_3_ x 6H_2_O, 0.41 mM CaCl_2_, 0.76 mM CoCl_2_, 0.66 mM ZnSO_4_ x 6H_2_O, 0.28 mM CuSO_4_, 0.19 mM Na_2_MoO_4_ x 2H_2_O, 0.38 mM NiCl_2_ x 6H_2_O and 0.19 mM VCl_3_. The final pH was adjusted to 6.0 by addition of NaOH pellets. The final media were subsequently made anoxic by several degassing and N_2_-addition cycles (minimum 25 cycles). The same medium, but without NH_4_Cl, was used for diazotrophic cultures. Na_2_S was used as both a reductant and sulfur source at a final concentration of 1.5 mM in all cases.

### Adaptation to diazotrophic conditions

*M. thermolithotrophicus* was adapted to growth under diazotrophic conditions after NH_4_Cl depletion from the media by three successive transfers to the same media without NH_4_Cl with a headspace containing 1.2 bar H_2_:CO_2_:N_2_ (58.2%:14.5%:27.3%) as described above. Cultures grown in media without NH_4_Cl with a gas phase of 1 bar H_2_:CO_2_ (80%:20%) were used as negative controls.

### Influence of trace metal availability on diazotrophic growth and tungstate inhibition

To determine influences of Mo and V on diazotrophic growth, we depleted media of Mo and V by three successive transfers of diazotrophic *M. thermolithotrophicus* cultures to media prepared as described above, but without Na_2_MoO_4_ x 2H_2_O or VCl_3_ or without both. The minimal concentration of Mo for diazotrophic growth was determined in a series of incubations, in which an already Mo-depleted diazotrophic culture was supplemented with Na_2_MoO_4_ x 2H_2_O concentrations ranging from 0.01 to 100 μM. W inhibition was tested by supplementing the already W-depleted diazotrophic culture with 0.001 to 10 μM Na_2_WO_4_ x 2H_2_O. A supplementation with the concentration of 100 μM gave unreproducible results (data not shown). W depletion was done as described above, with transfers to the media prepared without Na_2_WO_4_ x 2H_2_O.

### Cultivation in a fermenter

*M. thermolithotrophicus* was continuously grown in a 10 L fermenter (BIOSTAT^®^ B plus, Sartorius, Göttingen, Germany) under diazotrophic conditions with either 10 mM Na_2_SO_3_ or Na_2_SO_4_ as sulfur source instead of Na_2_S. The final culture volume was 7 L for the culture grown with Na_2_SO_3_ and 6 L for the culture grown with Na_2_SO_4_. The culture was continuously sparged with H_2_:CO_2_ (80%:20%) and N_2_ in the ratio of 1:1 and stirred with the speed of 500 rotation per minute, at 65 °C. As an inoculum, cultures cultivated in the same media were used in a ratio of 1:10.

### Ammonia measurement

Ammonia concentrations were measured in the culture supernatant: 0.5 mL culture aliquots were subsampled aerobically at each time point, cells were pelleted by centrifugation for 5 min at 15,700 x *g* using 5415R Microcentrifuge (Eppendorf, Hamburg, Germany) and the supernatant was frozen at −20° C until further use. Ammonia was measured by the salicylate-nitroprussidine method (90) in ROTILABO^®^ F-profile microtitration plates (Carl Roth GmbH, Karlsruhe, Germany). Standards in the range from 0 to 600 μM NH_4_Cl were prepared in the same medium used for cultivation. 80 μM of salicylate reagent (424.7 mM sodium salicylate, 193.8 mM tri-sodium citrate dihydrate, 193.8 mM di-sodium tartrate dihydrate and 0.95 mM sodium nitroprusside dihydrate) and 80 μM of hypochlorite reagent (10% sodium hypochlorite and 1.5 M NaOH mixed in the ratio of 1:36) were added to 40 μl of each sample. The plate was additionally mixed for 5 minutes on a shaker and incubated in the dark at the room temperature for 45 min. The absorbance was measured at 650 nm using an Infinite 200 PRO plate reader (Tecan, Männedorf, Switzerland) at room temperature.

### Evolutionary analyses

NifHDK, VnfHDK and AnfHDK sequences of 35 selected species were aligned using MUSCLE (91) (default parameters) in MEGA11 (92, 93). Afterwards, ambiguous positions were removed for each possible pairing (pairwise deletion option). The final alignment contained a total of 485 NifH positions, 729 NifD positions and 563 NifK positions. The sequence’s evolutionary history was inferred using the Neighbor-Joining method (94) with the JTT matrix model for multiple substitutions (95). The analysis was conducted in MEGA11 with ChlLNB from *Chlorobium limicola* as outgroup.

### Transcriptomics set-up

The cultures for transcriptomic profiling were grown in triplicates in batch as described above, but scaled up to culture volumes of 60 mL in 1 L pressure resistant Duran bottles. Inocula used to start the cultures were adapted to the respective conditions prior to inoculation as described above. Subsamples were taken anaerobically on ice after 3 h, 21 h and 25 h, and transferred to an anaerobic chamber, where they were pelleted and subsequently frozen in liquid nitrogen immediately after being taken out of the anaerobic chamber. Samples were stored at −80 °C until further use. Samples were shipped on dry ice to the Max Planck Genome Centre in Cologne, where they were processed and sequenced. RNA extraction and quality control, including rRNA removal, were also done at the Max Planck Genome Centre in Cologne.

### Transcriptome sequencing and analysis

Transcriptome sequencing was performed on an Illumina HiSeq 3000 (San Diego, CA, USA). Information on the raw reads are summarized in Supplementary Table 1. Raw RNA reads were quality trimmed and repaired using the *bbduk* and *repair.sh* scripts of the BBMap v35.14 suite (https://sourceforge.net/projects/bbmap/). Reads with a minimum length of 70 bp and a quality score of 20 were filtered for ribosomal RNAs using SortMeRNA v3.0.3 (96). Remaining mRNA reads were mapped against the *M. thermolithotrophicus* reference genome using Bowtie2 (97) as part of the SqueezeMeta v1.3.1 pipeline (98). The *DESeq2* R package (99) was subsequently used to calculate log_2_-fold changes, standard errors, test statistics and adjusted p-values. Changes in expression levels with adjusted p-values (padj) <0.05 and a minimum twofold change ratio (log_2_-fold change of 1 or higher) were considered significant (54).

The high-quality genome of *M. thermolithotrophicus* was used as a reference for mapping RNA reads. The genomic DNA of *M. thermolithotrophicus* was isolated using the protocol from Platero *et al*. (100) and sequenced on PacBio Sequel II platform using a single SMRT cell at the Max Planck Genome Center in Cologne. The genome was assembled using Flye v2.7 (101) and annotated as part of part of SqueezeMeta pipeline. Briefly, the ORFs were predicted using Prodigal (102) and similarity searches for GenBank (103), eggNOG (104), KEGG (105), were done using Diamond (106). HMM homology searches were done by HMMER3 (107) for the Pfam database (108).

### Sequence Submission

The *M. thermolithotrophicus* genome sequence and the transcriptome raw reads are available under the ENA project number PRJEB53446.

### Structural modeling and bioinformatic analysis

Alphafold2 was run with default parameters for all generated models. Predictions of membrane regions and overall topology were run on the DeepTMHMM server (https://dtu.biolib.com/DeepTMHMM/)(109) as of June 20^th^, 2022).

## Acknowledgements

We thank the Max Planck Institute for Marine Microbiology for continuous support and the Max Planck Genome Centre Cologne for RNA library preparation and sequencing. We also thank Dr. Mark Schweizer for setup the Alphafold2 pipeline and Dr. Susanne Erdmann for her helpful discussion regarding the virus-like/mobile elements encoding regions.

## Author contributions

N.M. and T.W. designed the research. N.M. performed all culture experiments. C.S. processed the transcriptomic data. C.S., N.M. and T.W. interpreted the data and all authors wrote the paper.

