## Supplementary material for "Transcriptomics sheds light on N_2_-fixation strategies employed by a thermophilic member of the *Methanococcales*"

### Supplemental Material

**Supplementary Table 1.** Mapping statistics**.**

**Supplementary Table 2.** Summary of up/downregulated genes pairwise.

**Supplementary Table 3.** Upregulated genes coding for putative virus-like elements in region 1 and their conservation among selected methanogens.

**Supplementary Table 4.** Downregulated genes coding for putative virus-like elements in region 2 and their conservation among selected methanogens.

**Supplementary Table 5.** Accession numbers for sequences used in phylogenetic reconstruction.

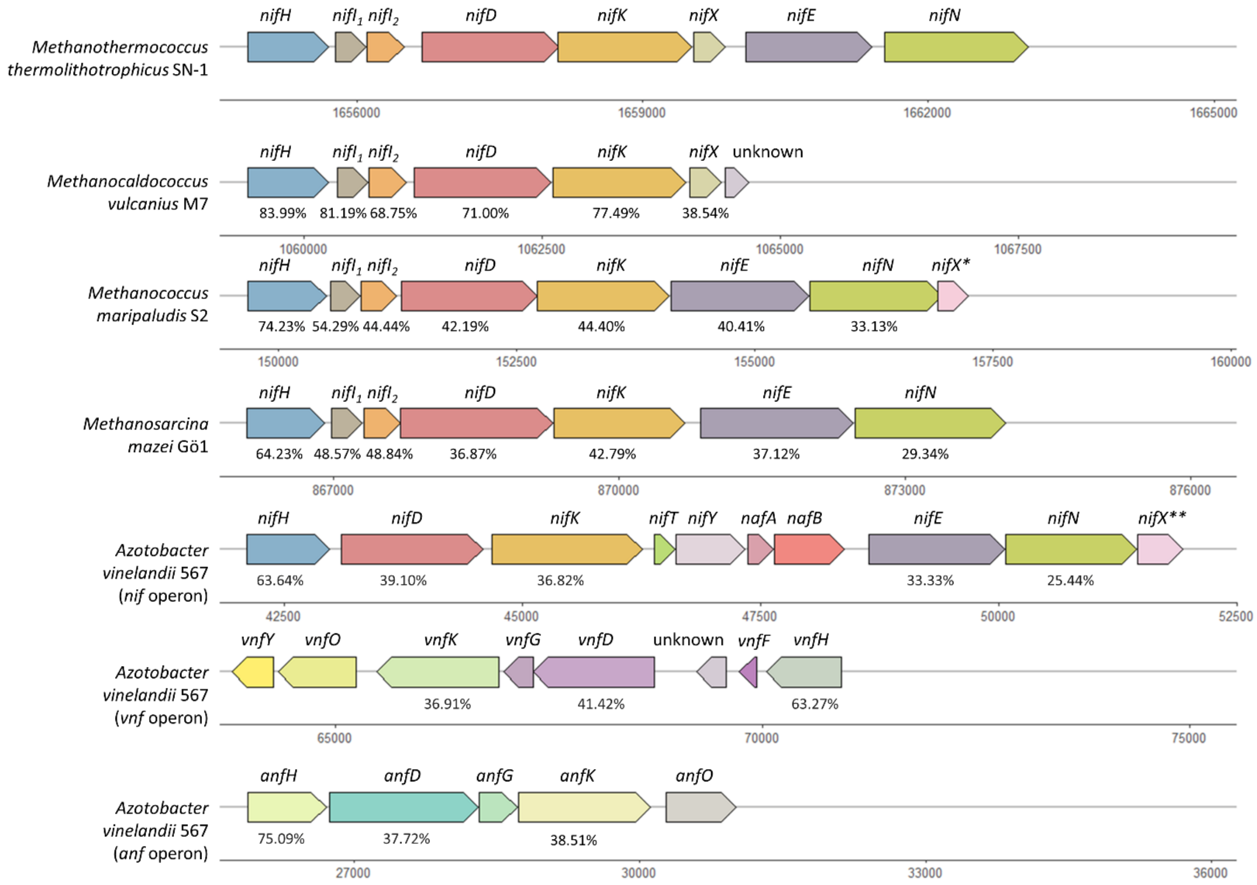


**FIG. S1** Genomic environment of *nif*, *vnf* and *anf* genes from selected diazotrophic methanogens and *A. vinelandii*. The gene *nifX* from *M. thermolithotrophicus* SN-1 (also referred as strain DSM 2095 in the main text), *nifX* from *M. maripaludis* S2 and *nifX* from *A. vinelandii* 567 do not share sequence homologies.

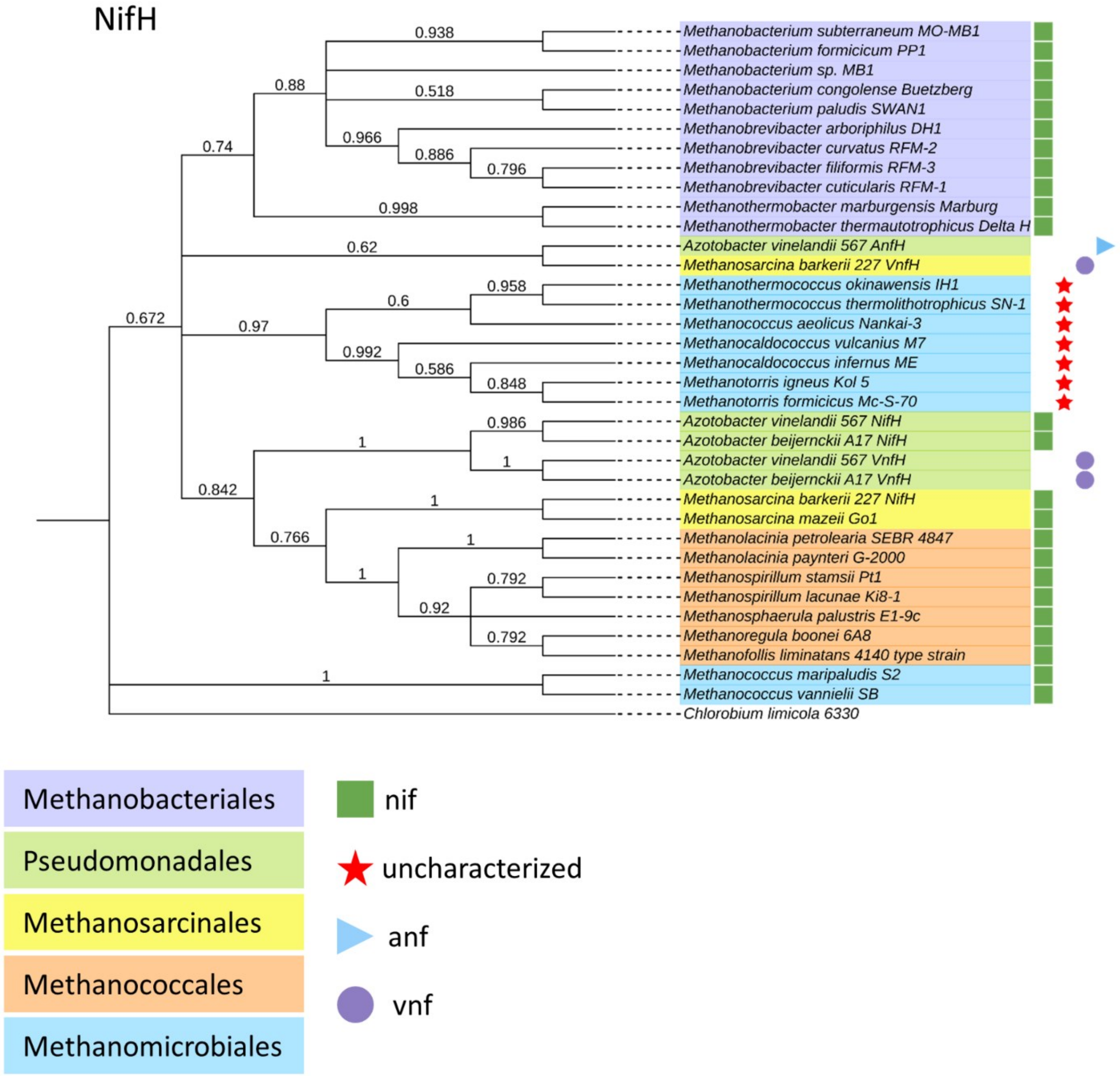


**FIG. S2** Evolutionary analysis of 35 NifH sequences. The percentage of replicate trees in which the associated taxa clustered together in the bootstrap test (500 replicates) are shown next to the branches (1). Evolutionary distances (2) are in the units of the number of amino acid substitutions per site. ChlL (light-independent protochlorophyllide reductase) from *Chlorobium limicola* was used as an outgroup. Accession numbers for sequences used in phylogenetic reconstruction can be found in Supplementary Table 5.

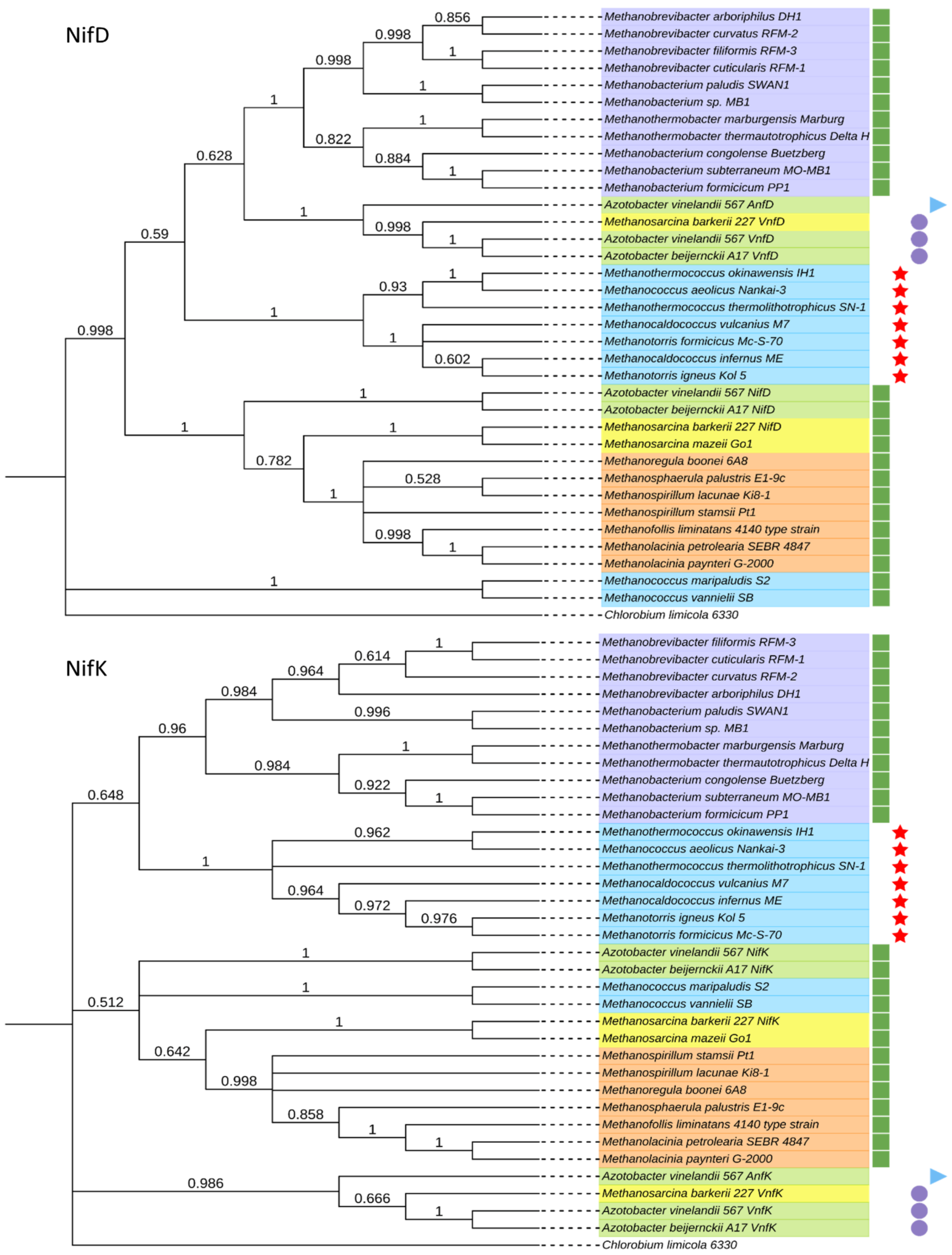


**FIG. S3** Evolutionary analysis of 35 NifD and NifK sequences. The percentage of replicate trees in which the associated taxa clustered together in the bootstrap test (500 replicates) are shown next to the branches (1). Evolutionary distances (2) are in the units of the number of amino acid substitutions per site. ChlNB from *Chlorobium limicola* were used as an outgroup. The same color-coding and symbols are used as in Fig. S2. Accession numbers for sequences used in phylogenetic reconstruction are provided in Supplementary Table 5.

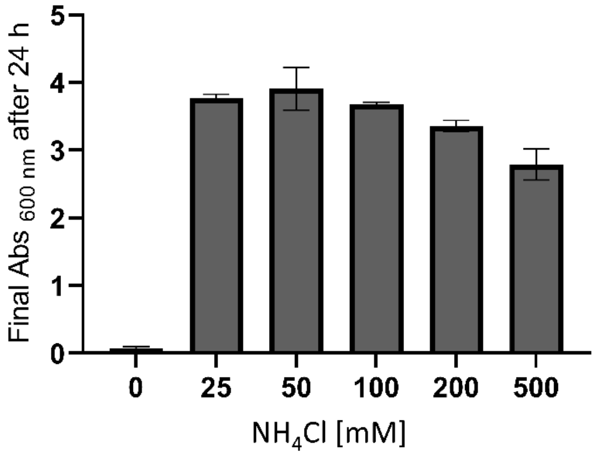


**FIG. S4** Final Abs $_{600nm}$ of *M. thermolithotrophicus* after 24 h of incubation with different $NH_4Cl$ concentrations. All measurements were done in triplicates.

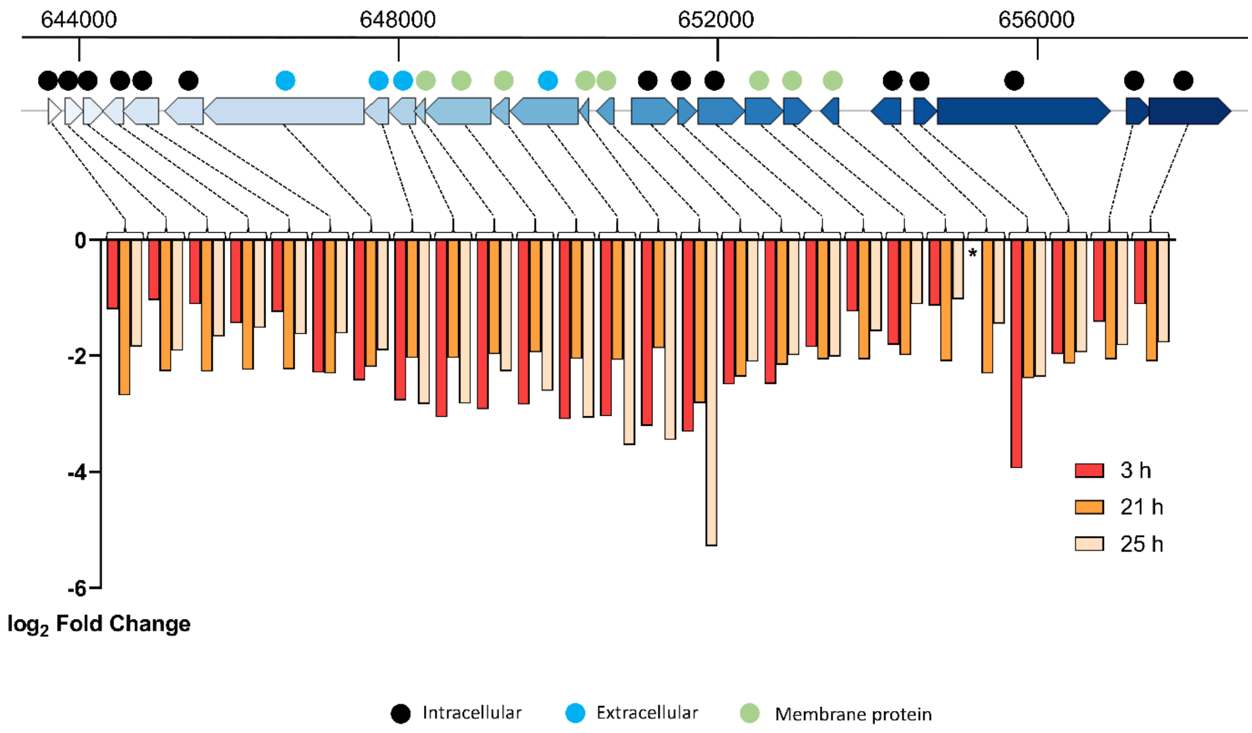


**FIG. S5** Log$_2$ fold changes overtime for the 26 co-occurring genes of unknown function. The downregulation of this genomic region in coordination with the upregulation of the other detected putative region of viral origin (Figure 5), might imply the competition of the two for the availability of transcriptional and translational machineries or is the result of a general cellular stress. The topology prediction was made via the DeepTMHMM server and is color-coded as follows: black for intracellular proteins, cyan for secreted proteins, and green for proteins containing transmembrane segments. WP numbers and predicted functions can be found in Supplementary Table 4. Gene that was not differentially expressed at 3 h is marked by an asterisk.
